# Fast Adversarial Generation of Molecular Dynamics Trajectories with Kinetic Fidelity

**DOI:** 10.64898/2026.01.21.700960

**Authors:** Nagesh B E, Subinoy Adhikari, Jagannath Mondal

**Affiliations:** Tata Institute of Fundamental Research, 36/P, Gopanpally Village, Serilingampally Mandal, Ranga Reddy District, Hyderabad 500046, Telangana, India

## Abstract

Molecular dynamics (MD) simulations yield atomic-level insights into molecular motion but struggle to reach the long timescales needed for rare events due to prohibitive computational costs. Generative machine-learning models (e.g., diffusion models and normalizing flows) offer a promising route to accelerate sampling, yet they generate independent equilibrium snapshots without temporal correlation or kinetic information. Autoregressive sequence models can learn time evolution by producing one frame at a time, but this stepwise generation often accumulates errors and drifts from true dynamics. Here, we propose a complementary approach inspired by advances in image and video generation: we treat finite MD trajectory segments as high-dimensional objects and learn their joint distribution using Generative Adversarial Networks (GANs). Using a Wasserstein GAN with gradient penalty, we directly generate entire time-series trajectories in one shot, that remain physically coherent over time without explicitly integrating the equations of motion. We demonstrate the generality of this method on molecular systems of increasing complexity: a 2D triple-well potential energy landscape, a protein–ligand binding process (cytochrome P450), the dynamics of an intrinsically disordered protein (*α*-synuclein) in a latent coordinate space, and even the conditional generation of folding trajectories for the Trp-cage mini-protein. In all cases, the GAN-generated trajectories closely reproduce the true free-energy landscapes and kinetic signatures of the systems, while enabling efficient sampling of rare events that would ordinarily require months of conventional MD simulation.

## INTRODUCTION

Understanding the time evolution of molecular systems via simulations requires solving Newton’s equations of motion. Molecular dynamics (MD) is a powerful tool for this task, providing detailed trajectories of atomic motions. However, conventional MD is computationally expensive and often fails to adequately sample the full phase space of complex systems. Rare events and long-timescale processes (e.g. protein folding) may require billions to trillions of simulation steps, far beyond practical limits [1]. This sampling bottleneck severely limits the applicability of MD to large systems or long-time predictions, motivating the development of new approaches to accelerate sampling [2].

In recent years, machine learning (ML) techniques, particularly generative models, have emerged as powerful tools for augmenting molecular dynamics (MD) simulations [3]. A central motivation for these efforts is the well-known limitation of MD in accessing long timescales and rare events, despite advances in hardware and enhanced-sampling algorithms. Generative ML models offer a complementary route: by learning from existing simulation data, they can potentially generate new configurations or trajectories at a fraction of the computational cost. A significant body of recent work has focused on static ensemble generation, where the primary goal is to reproduce the equilibrium Boltzmann distribution of molecular configurations. In this category, variational autoencoders (VAEs) [4], normalizing flows (e.g., Boltzmann Generators) [5], and diffusion probabilistic models [6] have demonstrated impressive success in generating unbiased molecular conformations and enhancing configurational sampling. Diffusion models, in particular, have been shown to efficiently explore complex free-energy landscapes and to reproduce equilibrium distributions with high fidelity [7, 8]. However, these approaches are fundamentally designed to sample from an explicitly learned probability distribution over configurations. As a result, individual samples are typically generated independently, and temporal correlations and kinetic information inherent to MD trajectories are not preserved by construction. While such models are invaluable for thermodynamic sampling, they are not naturally suited for generating continuous trajectories or reproducing time-dependent observables such as transition kinetics, dwell times, or correlated fluctuations.

A complementary line of research has therefore focused on learning the temporal structure of MD trajectories directly, often by treating discretized trajectories as sequences analogous to language. Sequential models such as Long Short-Term Memory (LSTM) networks [9] and, more recently, transformer-based GPT architectures [10], have been applied to learn state-to-state transitions and to predict the temporal evolution of molecular systems. These approaches can capture long-range dependencies and, in some cases, have demonstrated performance competitive with or superior to traditional Markov state models when predicting sequences of metastable states. Despite these successes, sequence-prediction models face important limitations when applied to MD. First, they are typically trained to predict the next step given previous context, making error accumulation unavoidable when generating long trajectories. Second, they often operate on discretized state representations, which can obscure fine-grained continuous dynamics. Finally, ensuring physical realism and global consistency over long timescales remains challenging, as local prediction accuracy does not guarantee that the generated trajectory obeys the correct joint statistics of MD.

These limitations highlight a key methodological gap: the need of generative models that can directly learn and sample from the joint distribution of entire time series, rather than from static ensembles or step-wise conditional distributions. In other scientific and engineering domains, this gap has been addressed using Generative Adversarial Networks (GANs) [11] for time-series generation. GAN-based frameworks have been successfully applied to produce realistic sequential data in diverse fields, including financial time series [12], physiological signals (e.g., ECG and EEG) [13], human motion capture [14], and audio generation [15]. Recurrent and time-aware GAN variants such as C-RNN-GAN, RGAN, and TimeGAN have explicitly demonstrated that adversarial learning can preserve temporal correlations autocorrelation structure, and event-time statistics in complex sequential data [16–18],. The common theme across these applications is that GANs learn an implicit joint distribution over entire sequences, allowing them to generate samples that are globally consistent in time rather than locally predicted step by step.

This property makes GANs a particularly natural candidate for MD trajectory generation. In contrast to explicit density models, a GAN does not require specifying or sampling from an explicit likelihood. Instead, the generator learns to produce entire trajectory segments that are indistinguishable from real MD trajectories under a discriminator (or critic) that evaluates the full time series. When the discriminator is exposed to complete trajectory windows, it can penalize violations of temporal smoothness, autocorrelation, and transition structure, thereby forcing the generator to reproduce these dynamical signatures implicitly. In this sense, GANs offer a direct route to learning the joint distribution of molecular configurations across time, which is precisely the object encoded by MD trajectories.

In this work, we leverage this perspective and employ Generative Adversarial Networks as trajectory-level generators for MD simulations. We treat finite MD trajectory windows as high-dimensional data samples, analogous to images or short video clips and train a GAN to learn their distribution. To ensure stable training and robust coverage of metastable states, we adopt the Wasserstein GAN with Gradient Penalty (WGAN-GP) formulation [19, 20]. WGAN-GP mitigates the well-known issues of mode collapse and training instability in vanilla GANs by optimizing the Wasserstein-1 distance between real and generated distributions while enforcing the Lipschitz constraint via a gradient penalty [20]. This formulation has proven effective in high-dimensional generative tasks, including time-series synthesis, where capturing subtle distributional differences is essential. By using WGAN-GP, our model learns the implicit joint distribution of consecutive MD frames and generates trajectory segments that preserve both equilibrium structure and temporal correlations. Unlike diffusion- or VAE-based approaches that generate independent configurations, and unlike autoregressive sequence models that predict trajectories step by step, the GAN framework produces entire time-series objects that are evaluated holistically. This enables faithful reproduction of dynamical signatures such as free-energy landscapes, transition kinetics, and implied timescales. As we demonstrate below across systems of increasing complexity, adversarial trajectory learning provides a natural and physically meaningful route for enhancing MD sampling and generating MD-consistent time series.

## RESULTS

### Representing MD trajectories as high-dimensional samples for adversarial trajectory synthesis

*a. Trajectory windows as samples.* A molecular dynamics (MD) trajectory is a multivariate time series in which the configuration at a given time is strongly correlated with its history. Our central idea is to cast *short trajectory windows* as individual high-dimensional samples, analogous to how images are treated as high-dimensional objects in computer vision (Fig. 1). In this representation, the *feature dimension* (e.g., *x/y/z* coordinates, or reduced/latent coordinates) plays the role of “channels”, while the time index plays the role of the spatial index in an image [11]. Learning the distribution over these trajectory-windows enables direct sampling of new time-consistent windows from a generative model.

**FIG. 1.**
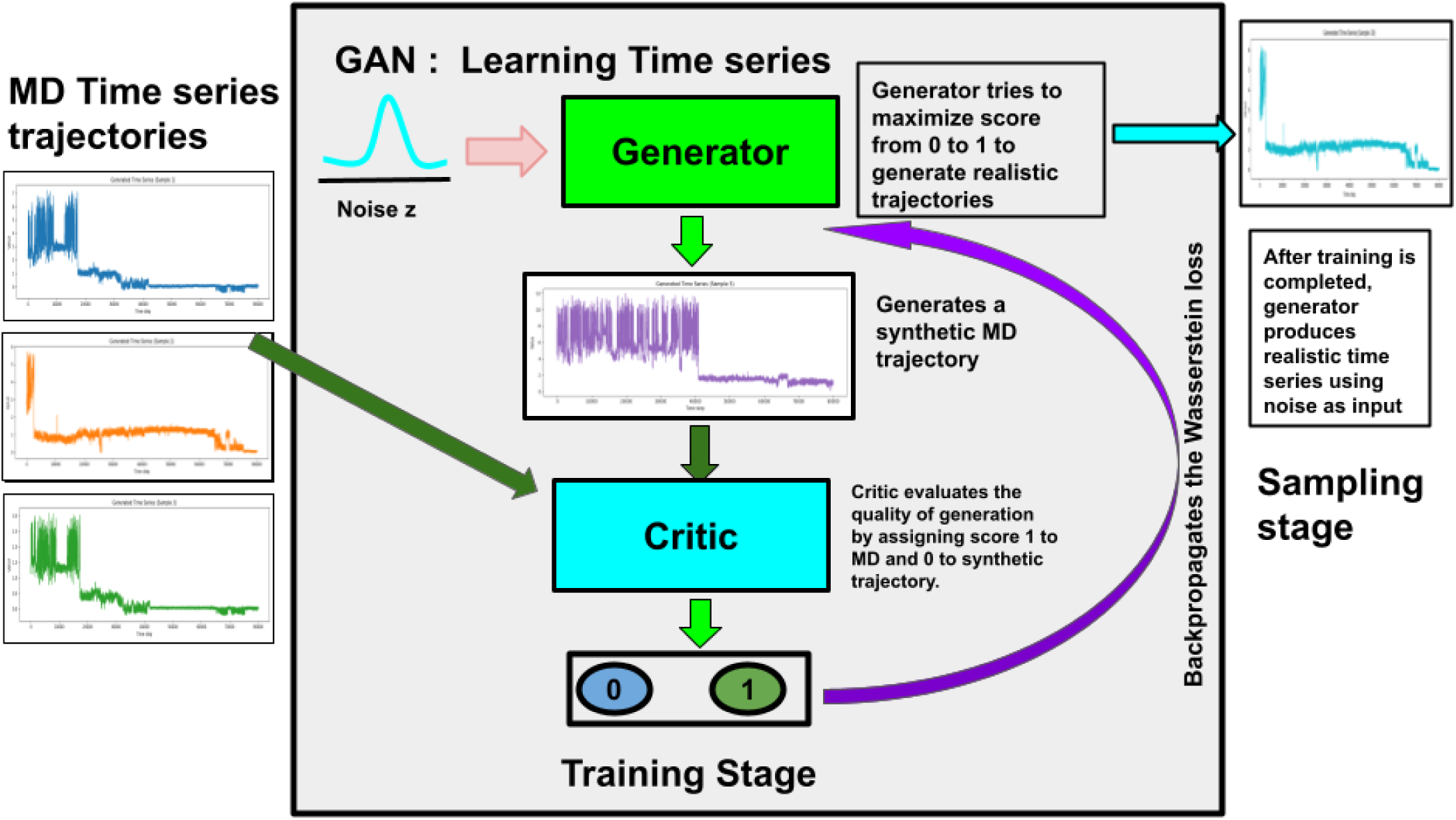
Workflow of the Model. We collected the MD simulation data from source simulations. Using random noise as input to generator, we generate synthetic trajectories. We pass both real and synthetic trajectories to a critic network which assigns the scores to real and fake as 1 and 0 respectively. During training stage, the generator tries to produce the trajectories which gets the score 1 from the critic. Critic becomes better at distinguishing the fake and real trajectories. Overall a adversarial optimization is done using Wasserstein’s loss function with gradient penalty for stable training. Once the training is done, we pass random noise to the generator to produce real trajectories which is the sampling stage.

Formally, let **x***_t_* ∈ R*^F^* denote an *F*-dimensional molecular descriptor at time *t* (e.g., *F* = 3*N* Cartesian coordinates, selected collective variables, or latent coordinates). A trajectory window of length *T* is

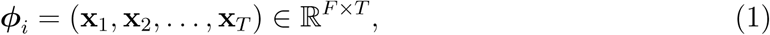

which we treat as the *i*-th training sample. This representation retains the ordering of time and therefore encodes temporal correlations implicitly through the joint structure of (**x**_1_*,…,* **x***_T_*).

*b. Why an adversarial model captures temporal structure.* Unlike models that generate independent frames, our generative model outputs an *entire trajectory window* ***φ*** at once. Consequently, the discriminator/critic evaluates realism using features that necessarily involve *relationships across time*—for example smoothness, lag-correlations, and transition-like patterns. Therefore, during training, windows that fail to reproduce temporal dependencies are assigned lower critic scores, forcing the generator to correct these deficiencies. In this sense, temporal information is captured because the model learns the *joint distribution P* (**x**_1_*,…,* **x***_T_*) rather than a set of independent marginals.

*c. Wasserstein GAN with gradient penalty (WGAN-GP).* We employ WGAN-GP, which improves stability and alleviates mode collapse by optimizing the Wasserstein-1 distance with a gradient penalty enforcing the 1-Lipschitz constraint [20]. The model consists of (i) a generator *G_θ_* that maps latent noise to a synthetic trajectory window, and (ii) a critic *D_ω_* that outputs a scalar score (not a probability).

**i) Generator.** The generator maps *z* ∼ *p_z_* (Gaussian noise in R*^dz^*) to a trajectory window ***ϕ̃*** = *G_θ_*(*z*) ∈ R*^FT^*. In our implementation, *G_θ_* is a fully-connected network with ReLU nonlinearities and an output layer of size *FT* reshaped to (*F, T*):

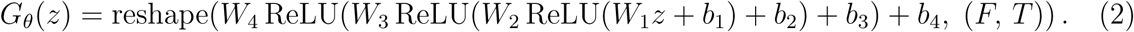

**ii) Critic.** The critic takes a trajectory window ***φ*** ∈ R*^F^* ^×*T*^ and outputs a scalar score

*D_ω_*(***φ***) ∈ R. We flatten ***φ*** into ***ϕ̃*** ∈ R*^F^ ^T^* and pass it through fully-connected layers with LeakyReLU activations:

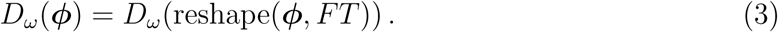

Higher scores correspond to samples that resemble the real trajectory-window distribution.

*d. Training objective.* Let ***φ*** ∼ *p*_data_ denote real trajectory windows and *z* ∼ *p_z_* latent noise. The WGAN-GP critic loss is

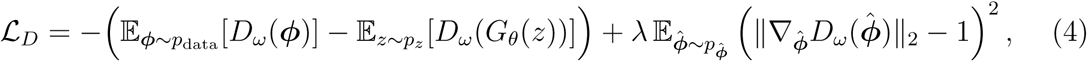

where ϕ̂ are linear interpolants between real and generated samples and *λ* is the gradient-penalty coefficient. The generator is trained to increase the critic score of generated samples:

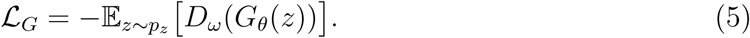

*e. Sampling and dynamical validation.* After training, we draw *z* ∼ *p_z_*and generate synthetic trajectory windows ***ϕ̃*** = *G_θ_*(*z*). Because these outputs are full time-series objects, we validate them using both thermodynamic and kinetic criteria. In subsequent sections, we quantify fidelity via free-energy surfaces in reduced coordinates and through kinetic diagnostics (e.g., transition statistics and Markov state model implied timescales), thereby t.com/esting whether the generated trajectories preserve the essential dynamical signatures of the underlying MD simulations.

We provide mathematical details of the wGAN-GP model in method section. Also we provide the first priciples theory of GAN model in Supplementary Information.

The details of the implementation of the model and sampling scripts can be found at the following URL: https://github.com/nagesh123-geek/GAN_MD

### GAN-based trajectory learning across increasing molecular complexity

To assess the generality of adversarial trajectory learning for molecular dynamics (MD), we benchmark our framework across a hierarchy of systems with systematically increasing dynamical and molecular complexity. Starting from a low-dimensional toy model with analytically interpretable kinetics, we progress through protein–ligand binding dynamics and finally to conformational dynamics of folded and intrinsically disordered proteins. This progression allows us to evaluate whether GAN-generated trajectories reproduce not only equilibrium distributions but also the essential temporal and kinetic signatures characteristic of MD.

### Two-dimensional three-well model: validating dynamical fidelity

We begin by validating the GAN-based trajectory learning framework on a minimal yet nontrivial dynamical system: a two-dimensional Brownian dynamics model of a single particle evolving on a three-well potential. This system serves as a controlled testbed, as it exhibits clearly defined metastable states, well-characterized transition pathways, and Markovian kinetics that can be directly assessed.

The reference MD trajectory, simulated under this three well potential, is segmented into fixed-length windows, each treated as a high-dimensional sample encoding the joint distribution of the (*x, y*) coordinates over time. Owing to the low dimensionality of the system, no additional dimensionality reduction or latent embedding is required, allowing the WGAN-GP model to operate directly on Cartesian coordinates and to generate full (*x*(*t*)*, y*(*t*)) time series. Synthetic trajectory windows generated after training are analyzed using the same thermodynamic and kinetic diagnostics as applied to the reference data. Figure 2(b) shows the real and generated trajectories in two dimension namely x(t) and y(t) for the Brownian motion that is used for training. The generated trajectories follow the dynamics closely as observed in simulation which confirms that, the generator sampling respects the time correlation and evolution of the system.

**FIG. 2.**
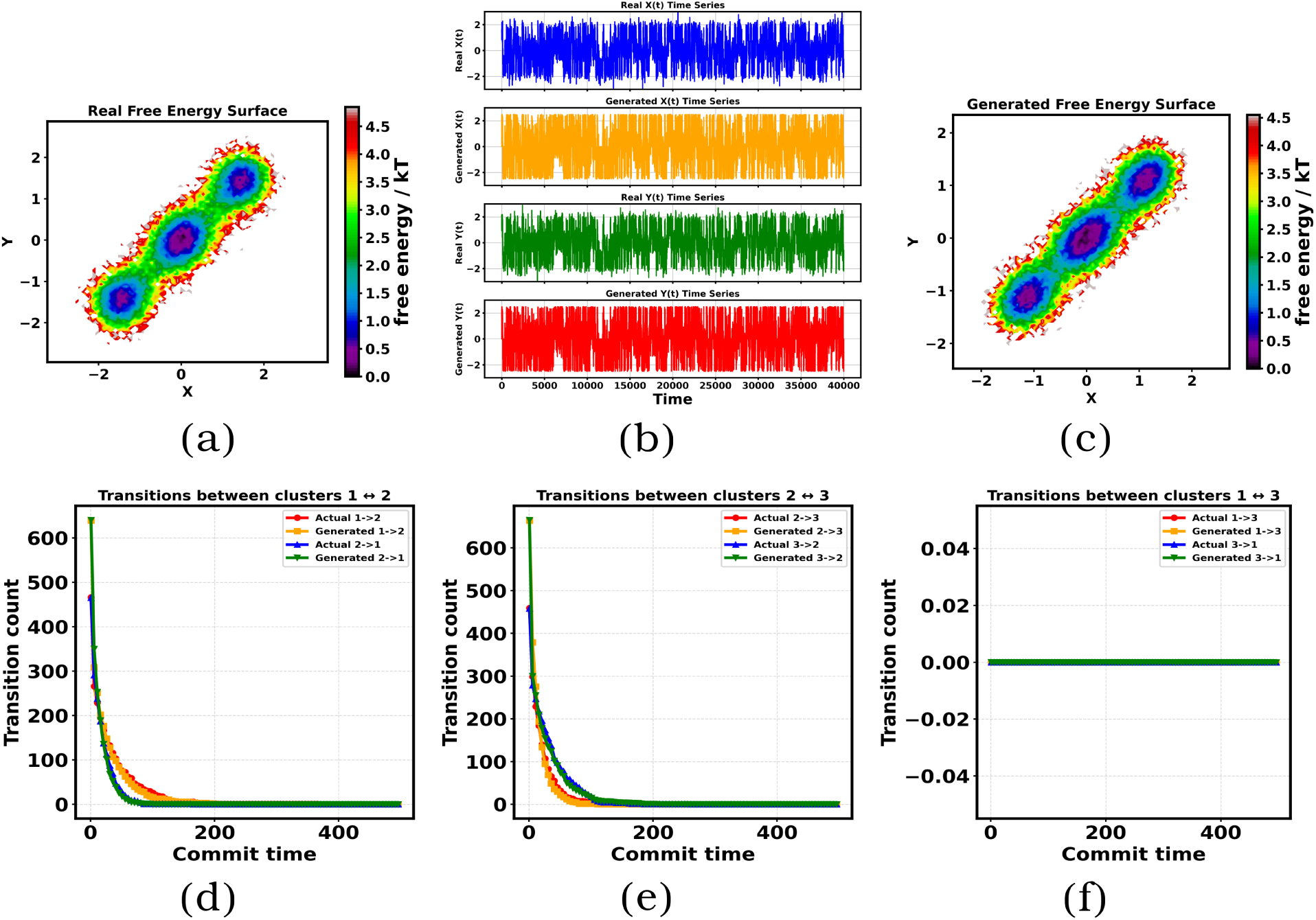
Dynamics of the two-dimensional three-well model. (a) Real free-energy surface (FES) of the three well system in the X-Y space. The particle transits over time from one minimum to another according to governing dynamical equations. (b) Real and generated trajectories of a paticle in two dimension for three well potential. The first and third panels corresponds to simulated time series trajectories X(t) and Y(t) respectively. The second and fourth panels correspond to generated trajectories of X(t) and Y(t) respectively from the model. (c) Generated Free energy surface of the trajectories generated using GAN in X-Y space. (d) State-to-state transition between minima 1 and 2 for chosen commit times. (e)State-to-state transition between minima 2 and 3. (f)State-to-state transition between minima 3 and 1.

Figures 2(a) and 2(c) compare the free-energy surfaces (FES) reconstructed from the real and GAN-generated trajectories, respectively. The generated FES faithfully reproduces the three metastable basins observed in the reference simulation, including their relative depths and spatial arrangement. Importantly, the connectivity between basins—manifested as continuous low-free-energy pathways—is also preserved, indicating that the GAN captures not only state populations but also the structural organization of the underlying energy landscape. This agreement suggests that the generator has learned the equilibrium distribution associated with the dynamics, despite being trained purely on finite trajectory windows.

To further probe dynamical fidelity, we assess whether the generated trajectories preserve the transition kinetics between metastable states. The trajectories are clustered into three states corresponding to the FES minima, and state-to-state transition statistics are computed as a function of lag (commitment) time. Figures 2(d–f) compare transition counts between all pairs of states for real and generated trajectories, including both forward and reverse transitions. The close agreement across lag times demonstrates that the GAN-generated trajectories reproduce the effective Markovian kinetics of the system. In particular, the decay of transition counts with increasing lag time follows the same trends in real and synthetic data, indicating that the generator captures the characteristic timescales associated with barrier crossings.

Taken together, the agreement at both the thermodynamic level (free-energy surface) and the kinetic level (state-to-state transition statistics) establishes that adversarial learning of trajectory windows can reproduce the essential dynamical signatures of this model system. Because the generator outputs full (*x, y*) time series, representative real and synthetic trajectories can also be visualized directly in configuration space, providing an intuitive illustration that the GAN produces physically continuous trajectories rather than disconnected samples. This low-dimensional example therefore provides a stringent and interpretable validation that WGAN-GP can learn the joint distribution of molecular coordinates across time, setting the stage for applications to more complex molecular systems.

### Protein–ligand binding dynamics: rare-event statistics in a biomolecular system

We next examine whether the GAN-based framework can be leveraged to synthesize protein–ligand binding trajectories, a class of rare-event dynamics that is notoriously challenging to capture using all-atom molecular dynamics (MD) [21, 22] Substrate recognition and binding in enzymes typically occur on microsecond to millisecond timescales, making direct observation via unbiased MD computationally expensive. As a representative and biologically relevant test case, we focus on substrate binding to cytochrome P450, a system for which even a single binding event requires multi-microsecond simulations.

In our previous MD studies of this system [23], capturing a single binding trajectory required approximately 3–4 weeks of wall-clock time on a GPU workstation using GRO-MACS, one of the fastest available MD engines. Due to this substantial computational cost, only three independent MD trajectories could be generated and analyzed. This severe sampling limitation motivates the use of generative approaches capable of producing additional statistically consistent trajectories at negligible computational cost.

To enhance sampling, we trained the GAN-based model on these three independent trajectories, using the time series of protein–ligand distance as a reduced yet physically meaningful observable. This distance coordinate provides a direct signature of binding and unbinding events, characterized by long dwell times in the unbound state punctuated by sharp transitions into the bound basin. The MD trajectories were segmented into fixed-length windows and used to train the WGAN-GP model, after which a large number of synthetic distance trajectories were generated. Binding events in both real and generated trajectories were identified using the same operational criterion (defined in the Methods section), ensuring a consistent kinetic comparison. Figure 3 summarizes the comparison between real and GAN-generated binding dynamics. Panel (b) shows the protein–ligand distance as a function of time for the three MD trajectories used for training. These trajectories illustrate the rarity of binding events and the long waiting times associated with substrate recognition. Panel (c) displays multiple GAN-generated distance trajectories, demonstrating that the model produces diverse realizations that exhibit similar qualitative features: extended unbound plateaus, sharp binding transitions, and bound-state residence times comparable to those observed in MD.

**FIG. 3.**
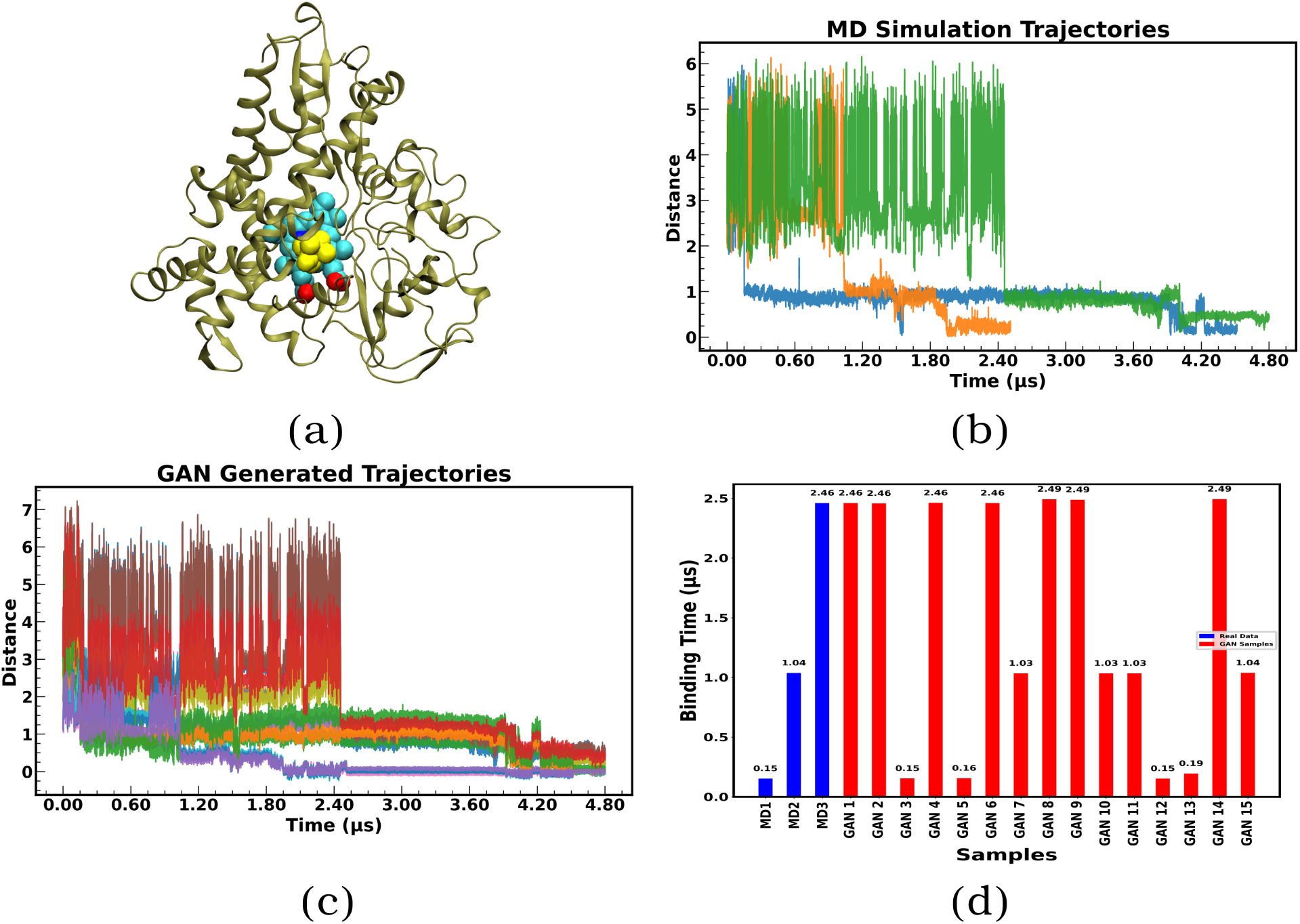
Protein–ligand binding dynamics in cytochrome P450. (a)A pictorial representation of Cytochrome P450 (b) Protein–ligand distance as a function of time for three independent MD trajectories used for training, illustrating rare binding events and long dwell times. (C) Representative GAN-generated distance trajectories showing similar binding–unbinding patterns and temporal variability. (d) Comparison of binding times extracted from real MD trajectories (blue) and GAN-generated trajectories (red), demonstrating quantitative agreement in rare-event statistics.

Importantly, the GAN framework delivers orders-of-magnitude acceleration over conventional MD. Training the model for 2000 epochs requires only 33.27 minutes. After training, loading the saved model and generating the first synthetic trajectory takes 4.6876 seconds, and subsequent trajectories can be produced in just 0.1487–0.2035 seconds per sample once the model is in memory. By contrast, generating comparable trajectories via unbiased all-atom MD on our available resources requires 30 days. Taken together, these timings highlight a significant speedup that enables rapid generation of statistically enriched trajectory ensembles, substantially improving sampling and rare-event statistics at a fraction of the computational cost.

Panel (d) compares the binding times extracted from the real MD trajectories (blue bars) with those obtained from multiple GAN-generated samples (red bars). The close correspondence between the two distributions indicates that the GAN captures not only equilibrium fluctuations in the distance coordinate, but also the statistics of rare, temporally extended binding events. Because binding times are highly sensitive to long-range temporal correlations and barrier-crossing kinetics, this agreement provides strong evidence that adversarial learning preserves essential dynamical information rather than merely reproducing marginal distributions. Taken together, these results highlight a key advantage of the GAN-based approach: once trained on a limited set of expensive MD trajectories, the model can generate a large ensemble of statistically consistent binding trajectories at negligible additional cost. This capability effectively amortizes the computational expense of rare-event MD simulations and enables rapid exploration of binding-time statistics that would otherwise require weeks to months of additional simulation.

### Intrinsically disordered protein *α*-synuclein: dynamics on a rugged landscape

As a more challenging test, we apply the framework to the intrinsically disordered protein *α*-synuclein, whose conformational dynamics span a broad and rugged free-energy landscape. To focus on slow collective motions, we analyze MD-simulated trajectories in a two-dimensional latent space obtained from a previously trained *β*-VAE [24] as shown in Figure 4(a). GAN-generated latent trajectories are compared directly to the reference MD-simulated latent trajectories. Figures 4(b) shows the latent trajectories from VAE model. Figure 4(c) shows the generated trajectories from GAN model after training with latent data.

**FIG. 4.**
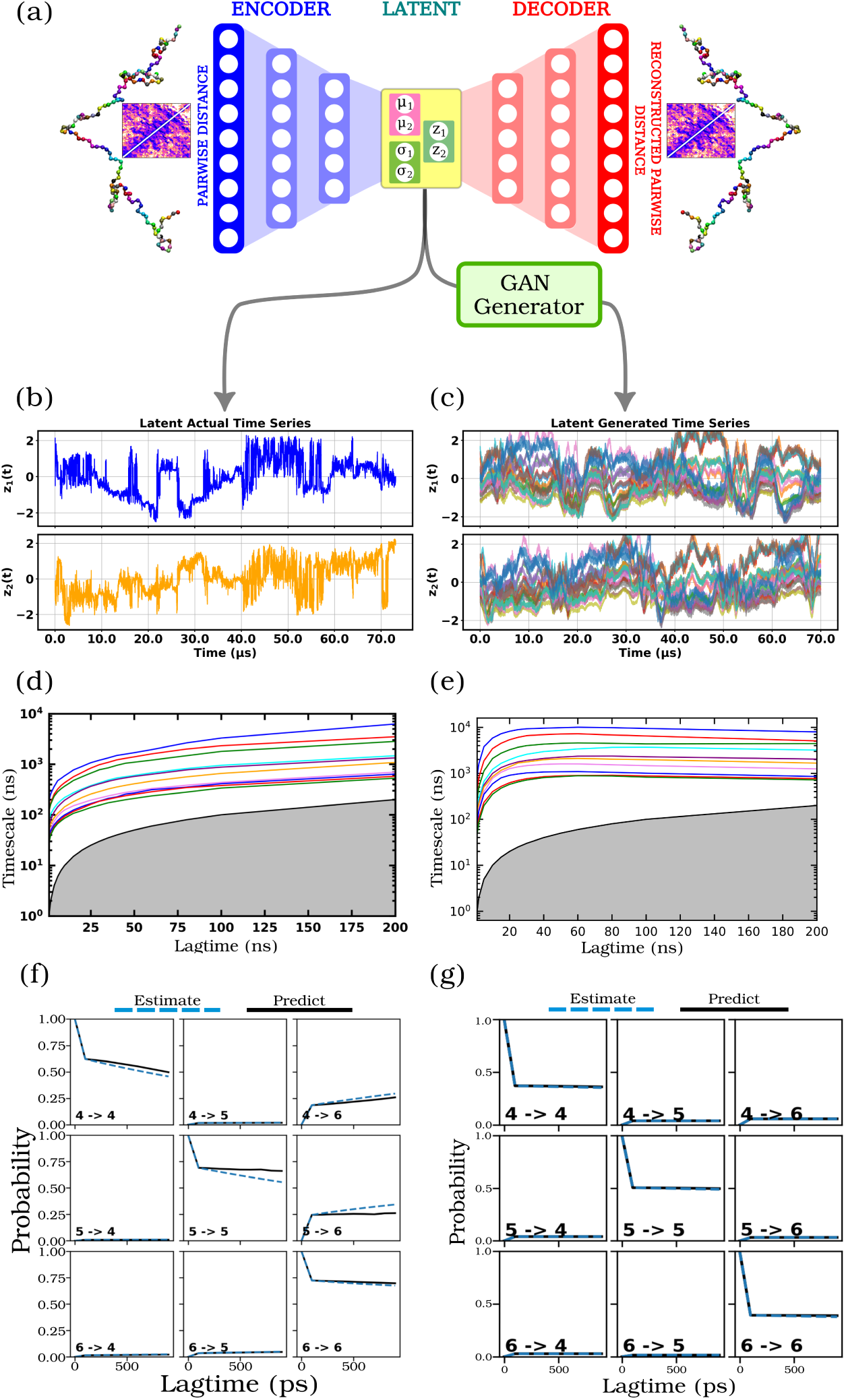
*α*-synuclein dynamics in latent space. (a)VAE architecture used to reduce high dimensional contact map data to two-dimensional latent data which is used for GAN training.(b) VAE-reduced two-dimensional latent dynamics *Z*_1_(*t*) and *Z*_2_(*t*) respectively. (c) GAN generated latent dynamics after training using Generator network with guassian noise as input for *Z*_1_(*t*) and *Z*_2_(*t*) respectively. (d) Implied time scales from MSM analysis for VAE reduced latent dynamics(e) Implied Time Scales for GAN generated latent dynamics. CK-test for selected states for (f) real data (g) generated data.

The kinetic consistency of the generated trajectories is assessed using Markov State Model (MSM) analysis. Figures 4(d,e) show implied timescale (ITS) plots obtained from real and generated trajectories, respectively. The agreement between the two indicates that GAN-generated trajectories capture the slow dynamical processes governing *α*-synuclein conformational rearrangements, despite the high ruggedness of the underlying landscape. The generated ITS Figures 4(e) exhibits improved convergence and smoother plateau behavior compared to the real ITS Figures 4(d). This result suggests that the GAN-generated data captures the underlying slow dynamical processes by implicitly learning the trajectory distributions. The ITS values for the GAN were higher than those for the VAE, indicating that the GAN captures slower relaxation processes more effectively. This suggests that the GAN’s implicit learning ability enhances the model’s ability to identify transitions between more kinetically distinct or stable states. To further validate the kinetic fidelity of the generated trajectories, we performed the Chapman-Kolmogorov (CK) test on MSM constructed from both real and generated datasets. Figure 4(f)-(g) and Figure S1-S2 show the CK test results for the real and generated trajectories for a few states. For the generated trajectories, the predicted transition probabilities closely follow the directly estimated values across the tested lag times, demonstrating strong kinetic consistency. Importantly, the agreement observed for the generated data is comparable to, and in some cases smoother than, that obtained from the real trajectories. This indicates that the generative model preserves the temporal correlations necessary for accurate long-timescale kinetic propagation, despite being trained on finite-length trajectories.

The close agreement between CK predictions and estimated transition probabilities further supports the conclusion drawn from the implied timescale analysis: the generated trajectories faithfully reproduce the slow dynamical processes governing *α*-synuclein conformational rearrangements. Together, the ITS and CK test results demonstrate that the generative model does not merely reproduce static structural distributions, but also captures the essential kinetic pathways and relaxation processes underlying the system’s dynamics.

### Trp-cage mini-protein: conditional generation of extended trajectories

Finally, we turn to trajectory synthesis for the globular 20-residue Trp-cage mini-protein, whose folding–unfolding dynamics are organized hierarchically across timescales. A central technical bottleneck in learning such dynamics is that training a generative model directly on very long, contiguous trajectories is challenging: the generator must represent correlations over tens of thousands of frames, which (i) increases the effective sequence length the network must “remember”, (ii) amplifies GPU memory and optimization instability during training, and (iii) often yields samples that match *static* distributions while failing to preserve *long-lag* kinetic structure (e.g., implied timescales) when only short windows are modeled independently. These issues are particularly acute for Trp-cage, where small deviations in temporal statistics can strongly distort folding/unfolding kinetics.

To overcome this limitation and enable synthesis of extended trajectories, we adopt a conditional GAN (cGAN) formulation [25] in which trajectory windows are generated while conditioning on frame-level context information. We use 2D latent trajectories from our prior VAE-based analysis as training data [24]. Concretely, the latent time series of shape (*L,* 2) is partitioned into consecutive chunks of shape (*C,* 2) with *C* = 40000 (yielding 12 chunks in our case), and each chunk is assigned conditioning labels that encode its frame-level context, analogous to conditioning used in video-generation models. During inference, the generator receives both a noise vector and the conditioning labels, producing windowed samples that can be stitched to assemble longer, temporally coherent synthetic trajectories.

Conditioning provides an explicit “anchor” that ties each generated window to its intended temporal context, reducing drift across windows and stabilizing training without requiring the network to process the full length *L* at once. This is essential for kinetically sensitive systems: although short-window generation can reproduce marginal latent distributions, recovering reliable implied timescales typically requires long, continuous trajectories. By combining chunking with frame-level conditioning, we retain long-range kinetic structure while keeping model size and memory demands tractable, thereby enabling efficient synthesis of extended Trp-cage trajectories.

Figures 5(a,b) show that GAN-generated trajectories reproduce the principal basins of the Trp-cage free-energy landscape. Kinetic validation via implied timescale analysis (Figs. 5(c,d)) confirms that the generated trajectories exhibit Markovian behavior consistent with the reference MD data. This conditional generation strategy demonstrates a viable route toward hierarchical trajectory synthesis without explicit integration of equations of motion.

**FIG. 5.**
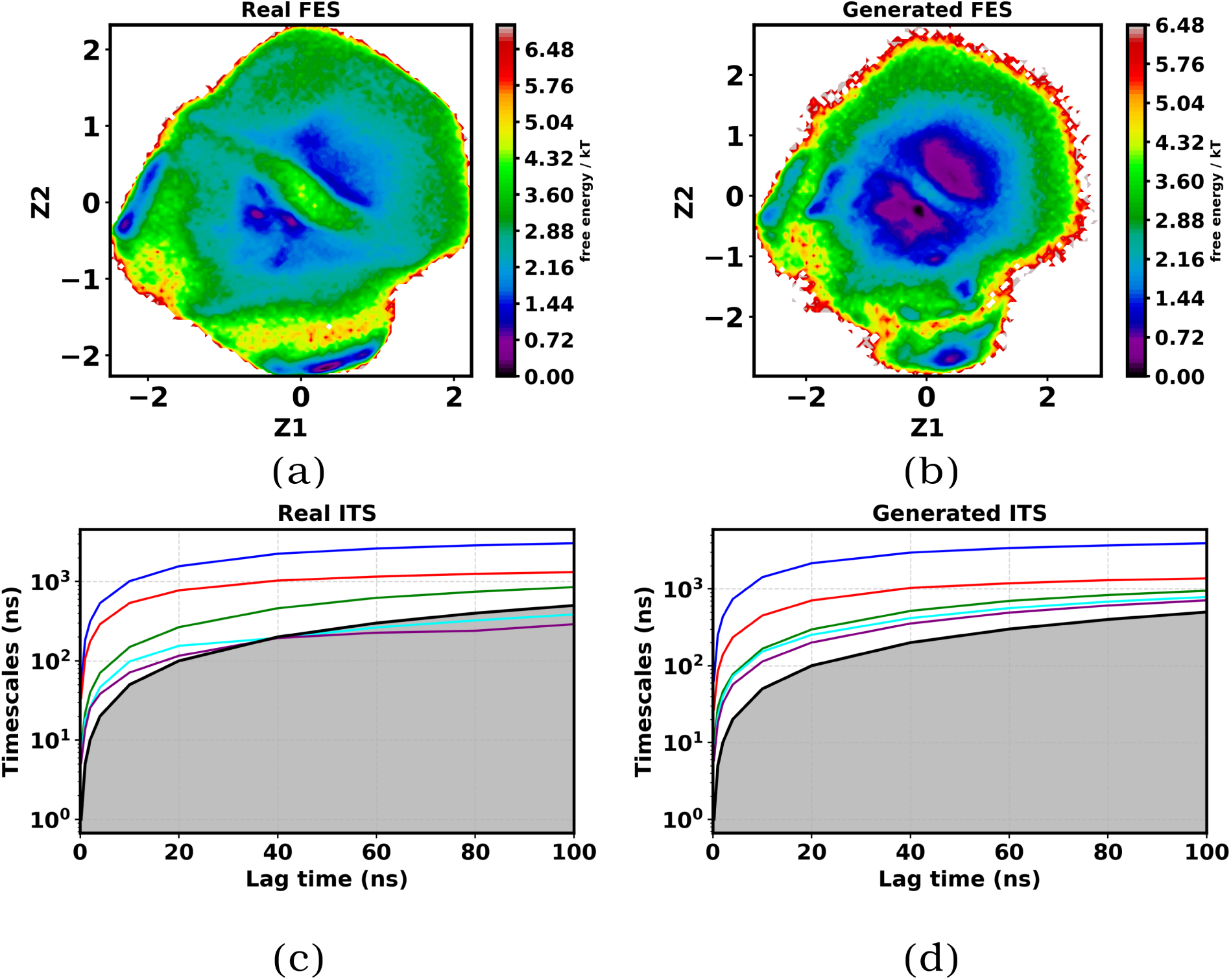
Trp-cage mini-protein. (a)Using VAE high dimensional contact map is reduced to two dimensional latent data. Figure shows the free energy surface in reduced *Z*_1_(*t*) and *Z*_2_(*t*) projectional coordinates. (b)Generated free energy surface using GAN after training the model with *Z*_1_(*t*) and *Z*_2_(*t*) latent coordinate data. (c) Implied time scale of VAE reduced latent *Z*_1_(*t*) and *Z*_2_(*t*). (d) Implied time scales of generated *Z*_1_(*t*) and *Z*_2_(*t*) from GAN after training.

## DISCUSSION

In this work, we repurpose adversarial generative modeling—originally developed for image and video synthesis—into a learn-the-whole-trajectory paradigm for molecular dynamics (MD). Rather than generating independent equilibrium snapshots or predicting frames autoregressively one step at a time, we treat short MD trajectory windows as high-dimensional objects and train a Wasserstein GAN with gradient penalty (WGAN-GP) to learn their joint distribution across time. In this trajectory-level setting, the critic evaluates an entire time series holistically, penalizing violations of temporal coherence, autocorrelation structure, and transition-like patterns. This adversarial objective therefore directly targets the statistics that define dynamics—free-energy landscapes (FES), transition pathways, and rare-event kinetics—without requiring explicit integration of equations of motion or likelihood-based modeling.

We evaluated this framework across a progression of systems with increasing physical and statistical complexity. In a two-dimensional three-well Brownian dynamics model, the generated trajectories reproduce both the equilibrium structure (FES) and kinetic observables, including state-to-state transition statistics, establishing that the adversarial objective captures meaningful temporal correlations beyond marginal distributions. We then moved to a rare-event regime by training on Cytochrome P450cam protein–ligand distance trajectories, where the GAN-generated time series recapitulate binding-time statistics consistent with unbiased MD, indicating that the model learns barrier-crossing dynamics and the temporal organization of recognition events. Next, for the intrinsically disordered protein *α*-synuclein, we assessed generated latent trajectories using Markov state modeling (MSM) diagnostics, finding that synthetic augmentation can improve the statistical robustness of kinetic estimates when MD data are limited. Finally, we addressed a practical bottleneck in trajectory generation: producing extended time series needed for kinetics-driven observables. By introducing frame-level conditioning—analogous to conditional video generation—we generated long Trp-cage trajectories whose implied timescales (ITS) and FES remain consistent with those from MD, supporting the view that conditional adversarial learning can maintain temporal coherence over long horizons.

Collectively, these four systems define a single conceptual axis of increasing difficulty, spanning (i) dimensionality, (ii) energetic ruggedness, (iii) rare-event kinetics and timescale separation, and (iv) hierarchical dynamics requiring long time-series synthesis. The three-well model provides an interpretable baseline where metastability and transitions can be validated directly. Protein–ligand binding introduces barrier-limited biomolecular events and strong timescale separation. Intrinsically disordered proteins add rugged landscapes and broad dynamical heterogeneity, making kinetics highly sensitive to finite sampling. Conditional Trp-cage synthesis targets the long-horizon regime, where reliable dynamical observables often demand continuous trajectories that cannot be inferred robustly from short windows alone. This systematic progression provides stringent evidence that trajectory-level adversarial learning can reproduce both thermodynamic and kinetic signatures of MD across distinct regimes.

These results complement and extend emerging efforts to generate MD time series using generative models. Prior work such as MD-GAN demonstrated physically consistent trajectory generation for relatively simple systems (harmonic oscillator, bulk water, polymer melts) [26]. In parallel, autoregressive sequence models—including LSTMs [9] and transformer-based architectures [10]—have been explored by discretizing dynamics into tokens and learning stepwise sequence rules. While effective in many settings, stepwise generation can accumulate errors over long horizons and does not guarantee global consistency of trajectory-level statistics. By contrast, the *learn-the-whole-trajectory* paradigm directly models the joint structure of trajectory windows: the generator proposes complete time-series objects, and the critic enforces realism using features that necessarily couple multiple time points. This makes adversarial learning a natural fit for trajectory realism—including long-lag kinetic diagnostics such as binding-time distributions, transition statistics, and implied timescales—rather than only for static ensemble reproduction.

Looking forward, an important opportunity is to move beyond system-specific generators toward models that learn *transferable* dynamical priors across proteins and sequence space. Achieving such transfer will likely require representations and objectives that disentangle “universal” dynamical motifs—diffusion along slow collective coordinates, barrier crossing, metastability—from system-dependent details such as force-field idiosyncrasies and specific interaction networks. One promising direction is the development of conditional foundation models for molecular time series, trained on diverse trajectory corpora and conditioned on descriptors such as sequence, structural context, thermodynamic state, or learned embeddings. Such models could support interpolation in sequence space, enabling rapid in silico screening and hypothesis generation for designed variants, while amortizing the cost of expensive MD campaigns across families of related systems.

Methodologically, hybrid objectives that combine adversarial learning with explicit temporal forecasting may further strengthen long-horizon consistency and move toward predictive dynamical surrogates. While our current framework can generate trajectories that match MD statistics when conditioned appropriately, it does not yet learn an explicit evolution operator that propagates a given initial condition forward in time. A natural next step is to integrate trajectory-level critics with autoregressive or state-space components (e.g., latent recurrent/transformer dynamics), where the generator is trained to both (i) satisfy a global realism objective over windows and (ii) obey multi-step consistency constraints that encourage faithful propagation of slow modes. Physics-informed regularization, such as constraints related to detailed balance in latent MSM space, consistency of drift–diffusion estimates, or stability constraints on learned propagators—could further improve interpretability and robustness.

Finally, expanding transferability requires rigorous benchmarking on out-of-distribution systems and sequence variants, using diagnostics that emphasize kinetics (implied timescales, committor structure, transition-path statistics) rather than only marginal distributions. If successful, transferable trajectory-level generators could serve as fast “trajectory amplifiers” that enrich rare-event statistics, guide adaptive sampling, and enable rapid hypothesis testing across protein families and engineered sequence landscapes. More broadly, the learn-the-whole-trajectory paradigm suggests a route for generative AI to transition from producing plausible short sequences to providing reusable, statistically grounded surrogates for complex dynamical processes in molecular science.

## METHOD

### GANs: A Special Case of Variational Divergence Minimization

Generative Adversarial Networks (GANs) can be viewed as a special case of the variational *f*-divergence framework corresponding to the *Jensen–Shannon divergence*.We refer SI for detailed mathematical tehory of GAN models in general.

#### Choice of f-divergence

For GANs, the function *f* (*u*) is chosen as:

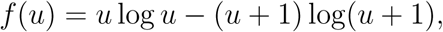

which corresponds to the Jensen–Shannon divergence (up to additive constants). The convex conjugate *f* ^∗^(*t*) and its domain are:

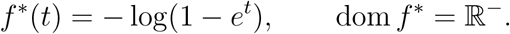

#### Critic Parameterization

To ensure *T_ω_*(*x*) ∈ dom(*f* ^∗^), the critic is parameterized as

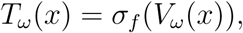

where

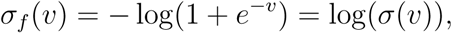

and 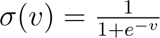 is the logistic sigmoid function. This ensures *T_ω_*(*x*) *<* 0 for all *x*.

#### Variational Objective

Substituting the chosen *f* ^∗^(*t*) into the general variational form gives

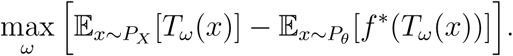

Expanding *f* ^∗^(*t*) explicitly,

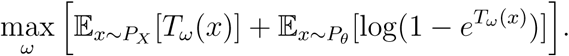

### Wasserstein GAN with Gradient Penalty (WGAN-GP)

The original WGAN enforced the 1-Lipschitz constraint on the critic *T_ω_*(*x*) by *weight clipping*, which often led to training instability and poor gradient propagation. To address this, Gulrajani et al. (2017) [20] introduced a smooth *gradient penalty* formulation, known as WGAN-GP.

#### Motivation

A function *T_ω_*(*x*) is 1-Lipschitz if and only if its gradient norm satisfies

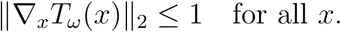

Instead of hard-clipping weights, WGAN-GP enforces this condition softly by adding a regularization term to the critic loss.

#### Gradient Penalty Term

The gradient penalty term is defined as:

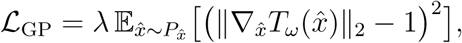

where

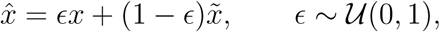

with *x* ∼ *P_X_* (real samples) and *x̃* ∼ *P_θ_* (generated samples). The distribution *P_x_*_^_ represents points uniformly sampled along straight lines between real and generated data points.

#### Final Objective Functions

The critic (discriminator) objective in WGAN-GP is:

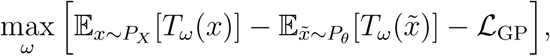

and the generator objective is simply:

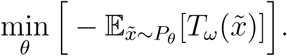

## NETWORK STRUCTURE AND HYPERPARAMETERS

We used pytorch [27] to implement GAN architecture. We used Adam optimizer [28]for optimization. We updated the critic for every fifth update of generator. Learning rate used is 0.00001. We below provide the architectural details for generator and critic neural networks.

*f. Generator.* The generator is a fully-connected neural network consisting of three hidden layers with ReLU activations and an output layer mapping to a signal of length *T* = signal length with *F* = n features. n represents actual MD trajectory dimension. For example for 2d three well potential we used, n = 2. Signal length is the total time frames in MD trajectory. Let *z* ∈ R*^dz^* be a latent noise vector, where *d_z_* = latent dimension.

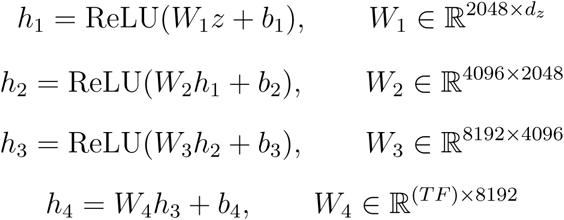

The final vector *h*_4_ ∈ R*^T^ ^F^* is reshaped into a two-dimensional output:

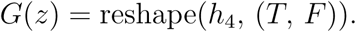

The generator can be expressed as:

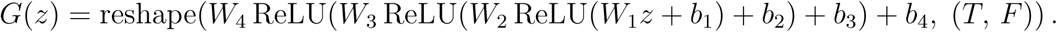

*g. Critic.* Let *x* ∈ R*^T^* ^×*F*^ be an input signal, where *T* = signal length and *F* = n features. The critic first flattens the input into a vector

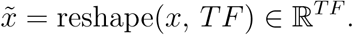

It then passes *x̃* through a sequence of fully-connected layers with LeakyReLU activations (slope 0.2):

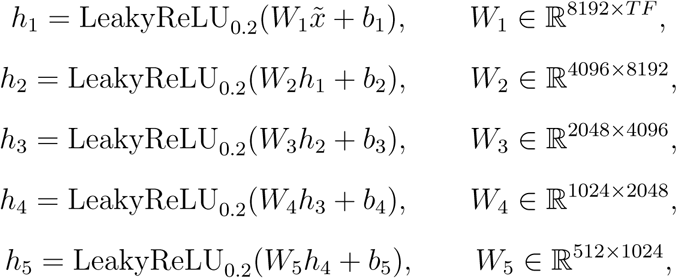

Finally, the critic outputs a scalar score:

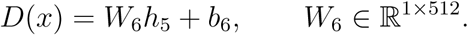

In compact form, the critic outputs a scalar score:

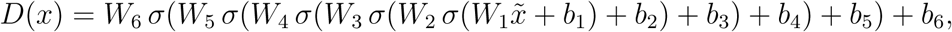

## DETAILS OF THE MODEL SYSTEMS

Here we provide the simulation details of the data used for training.

### Three well potential

The potential for the 3-state model is given by [9]

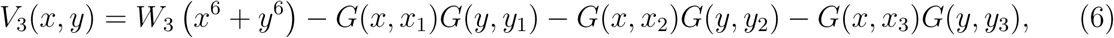

where *G* is a Gaussian function defined as

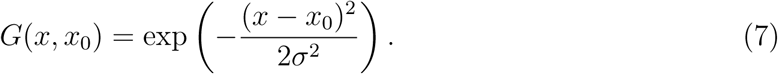

Here, *x*_0_ and *σ* denote the mean and standard deviation of the distribution, respectively. we set *W*_3_ = 0.0001 and *σ* = 0.8.

The mean values of the Gaussian distributions for the 3-state model are given by

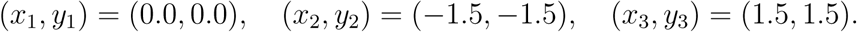

### Trp-cage mini protein and *α*-Synuclein

For Trp-cage and *α*-Synuclein, we used very long molecular dynamics simulation trajectories from D. E. Shaw Research (DESRES). The Trp-cage trajectory spanned 100 *µ*s, while the *α*-Synuclein trajectory was 73 *µ*s long. These simulations were performed using the a99SB-disp force field on Anton specialized hardware [29]. The detailed simulation protocols are found in the original paper by Robustelli et al. [30].

### Protein Ligand systems

The trajectories taken from the MD simulations are all not done to the same time. In order to be consitent with the dimension, we padded the zeros at the end to match the longest simulation length, as adding zeros at the end will not alter the dynamics and it is a standard practice in image analysis as in CNNs. Then we down sampled the entire trajectory to a length of 80000 by skipping 6 interval steps. The original simulation is done at 10ps, now it is reduced to 60ps.

## CODE AVAILABILITY

The details of the implementation of the model and sampling scripts can be found at the following URL: https://github.com/nagesh123-geek/GAN_MD

## SUPPLIMENTARY INFORMATION

The supplimentary information (SI) provides supplimentary figures and mathematical details of the model used. The SI section THEORY OF GAN MODELS discusses the GAN model from the f-divergence minimization perspective. We derive the objective function for the GAN using variational Lower Bound. Combining all the results at the end we derive the generalized f-GAN min max adversarial objective function on which GAN is optimized. Figure S1 provides the results of the CK-test for the real *α*-synuclein data. Figure S2 provides the results of the CK-test for the generated *α*-synuclein data.

## ACKNOWLEDGEMENT

We acknowledge support of the Department of Atomic Energy, Government of India, under Project Identification No. RTI 4007. All the authors acknowledge Tata Institute of Fundamental Research Hyderabad, India for providing the support of computing resources. JM acknowledges funding from Department of Science and Technology of India (CRG/2023/001426).

