## supplemental method and figures for "Fast Adversarial Generation of Molecular Dynamics Trajectories with Kinetic Fidelity"

(Dated: January 22, 2026)

*

†

#### THEORY OF GAN MODELS

#### GENERATIVE MODELLING VIA F-DIVERGENCE MINIMIZATION

Let the dataset be

$$D = \{x_1, x_2, \dots, x_n\} \sim P_X$$

where each data point $x_i \in \mathbb{R}^d$. The goal of generative modelling is:

1. Given data $D$, learn the underlying distribution $P_X$.

2. Once $P_X$ is approximated, generate (sample) new data points from it.

A general and powerful approach to this problem is to minimize a statistical divergence between the true data distribution $P_X$ and a parameterized model distribution $P_\theta$.

##### f-Divergence Definition

For two probability density functions $p_X$ and $p_\theta$, the *f-divergence* is defined as

$$D_f(P_X \| P_\theta) = \int p_\theta(x)\, f\left(\frac{p_X(x)}{p_\theta(x)}\right) dx$$

where $f : \mathbb{R}_+ \to \mathbb{R}$ is a convex, lower semi-continuous function satisfying $f(1) = 0$.

Different choices of $f(u)$ yield common divergence measures:

$$\begin{aligned} f(u) &= u \log u && \Rightarrow \text{Kullback–Leibler (KL) divergence,} \\ f(u) &= -\log u && \Rightarrow \text{Reverse KL divergence,} \\ f(u) &= (u-1)^2 && \Rightarrow \text{Pearson } \chi^2 \text{ divergence,} \\ f(u) &= |u-1| && \Rightarrow \text{Total Variation distance.} \end{aligned}$$

The optimization problem is then:

$$\theta^* = \arg\min_\theta D_f(P_X \| P_\theta)$$

##### Variational Representation via Convex Conjugate

To make $D_f$ computable, we use the Fenchel conjugate of $f$:

$$f^*(t) = \sup_{u \in \mathrm{dom}(f)} \big(ut - f(u)\big)$$

which implies

$$f(u) = \sup_{t \in \text{dom}(f^*)} \big(tu - f^*(t)\big).$$

Substituting into the definition of $D_f$:

$$\begin{aligned} D_f(P_X \| P_\theta) &= \int p_\theta(x) \sup_t \left[ t \, \frac{p_X(x)}{p_\theta(x)} - f^*(t) \right] dx \\ &= \sup_{T \in \mathcal{T}} \int p_\theta(x) \left[ T(x) \frac{p_X(x)}{p_\theta(x)} - f^*(T(x)) \right] dx \\ &= \sup_{T \in \mathcal{T}} \left( \mathbb{E}_{x \sim P_X}[T(x)] - \mathbb{E}_{x \sim P_\theta}[f^*(T(x))] \right) \end{aligned}$$

where $T : \chi \to \text{dom}(f^*)$ is a function (often a neural network) called the *critic* or *discriminator.*

**Variational Lower Bound**

Thus, we obtain the variational lower bound:

$$D_f(P_X \| P_\theta) \geq \sup_{T \in \mathcal{T}} \left( \mathbb{E}_{x \sim P_X}[T(x)] - \mathbb{E}_{x \sim P_\theta}[f^*(T(x))] \right)$$

**Parameterized Critic Function**

In practice, the supremum over the function space $\mathcal{T}$ is approximated by a neural network parameterized by weights $\omega$. Thus, we define the critic (or discriminator) as

$$T_\omega(x) = \sigma_f(V_\omega(x)),$$

where

- $V_\omega(x)$ denotes the pre-activation output of the neural network with parameters $\omega$,
- $\sigma_f(\cdot)$ is an activation function chosen to ensure that $T_\omega(x) \in \text{dom}(f^*)$, i.e., the range of $T_\omega(x)$ matches the domain required by the conjugate function $f^*(t)$.

This parameterization ensures that the critic's output remains in a valid range for each divergence. For instance:

$$f(u) = u \log u \quad \Rightarrow \quad f^*(t) = e^{t-1}, \quad \text{so } \sigma_f(\cdot) = \text{identity},$$

$$f(u) = -\log u \quad \Rightarrow \quad f^*(t) = -1 - \log(-t), \quad \text{so } \sigma_f(\cdot) = -\exp(\cdot),$$

$$f(u) = u \log u - (u+1)\log(u+1) + \log 4 \quad \Rightarrow \quad f^*(t) = -\log(1 - e^t) - \log 2, \quad \text{so } \sigma_f(\cdot) = \log \sigma(\cdot),$$

where $\sigma(\cdot)$ is the logistic sigmoid function.

**Final Objective with Parameterized Critic**

Substituting $T_\omega(x)$ into the variational bound, we obtain the practical training objective:

$$\min_\theta \max_\omega \left[\mathbb{E}_{x\sim P_X}[T_\omega(x)] - \mathbb{E}_{x\sim P_\theta}[f^*(T_\omega(x))]\right]. \tag{1}$$

This yields the general form of the *f-GAN* objective, where both the generator and critic are parameterized by deep neural networks.

### I. CK-TEST RESULTS OF $\alpha$-SYNUCLEIN

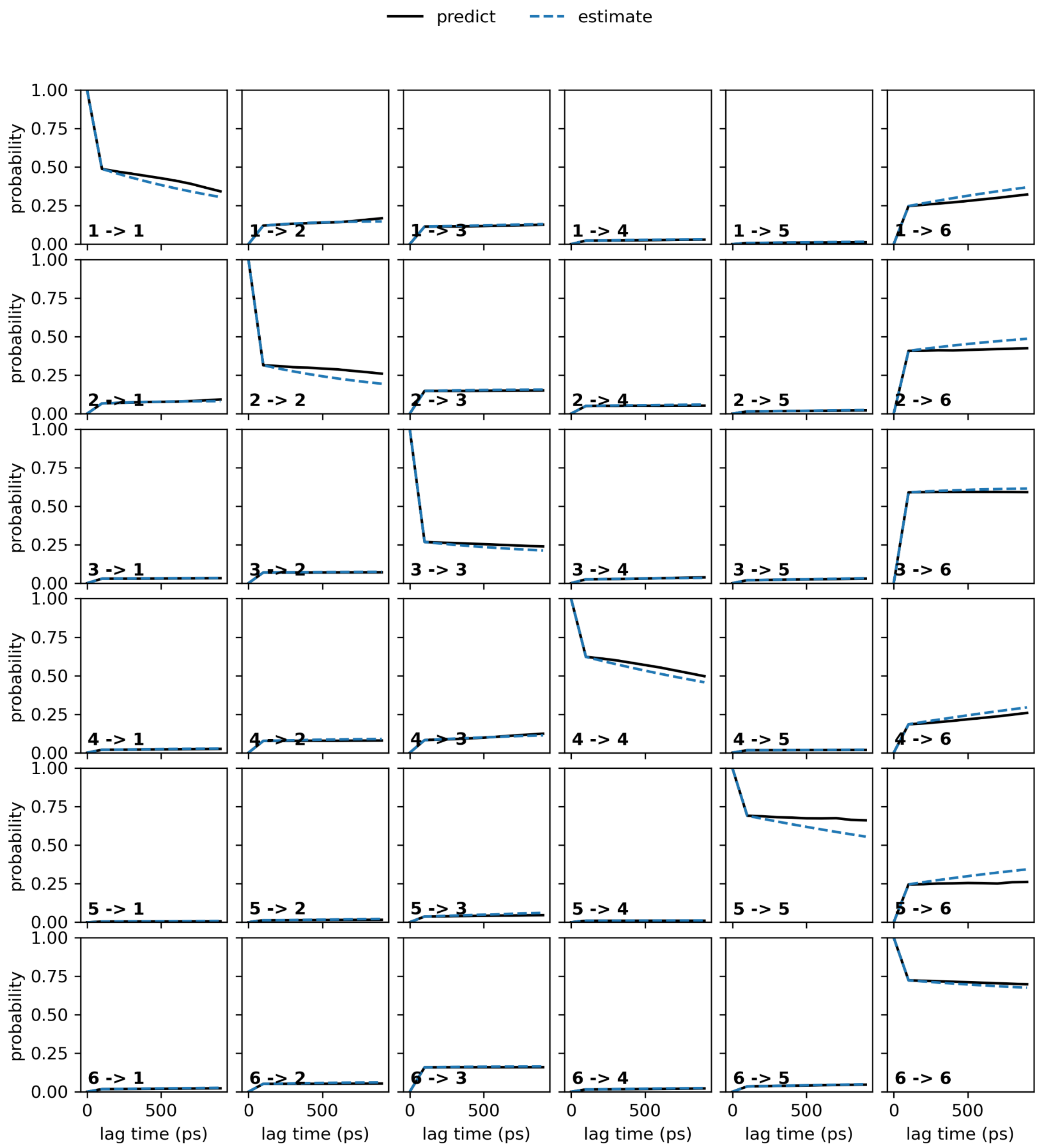


FIG. S1. CK-test for the real data.

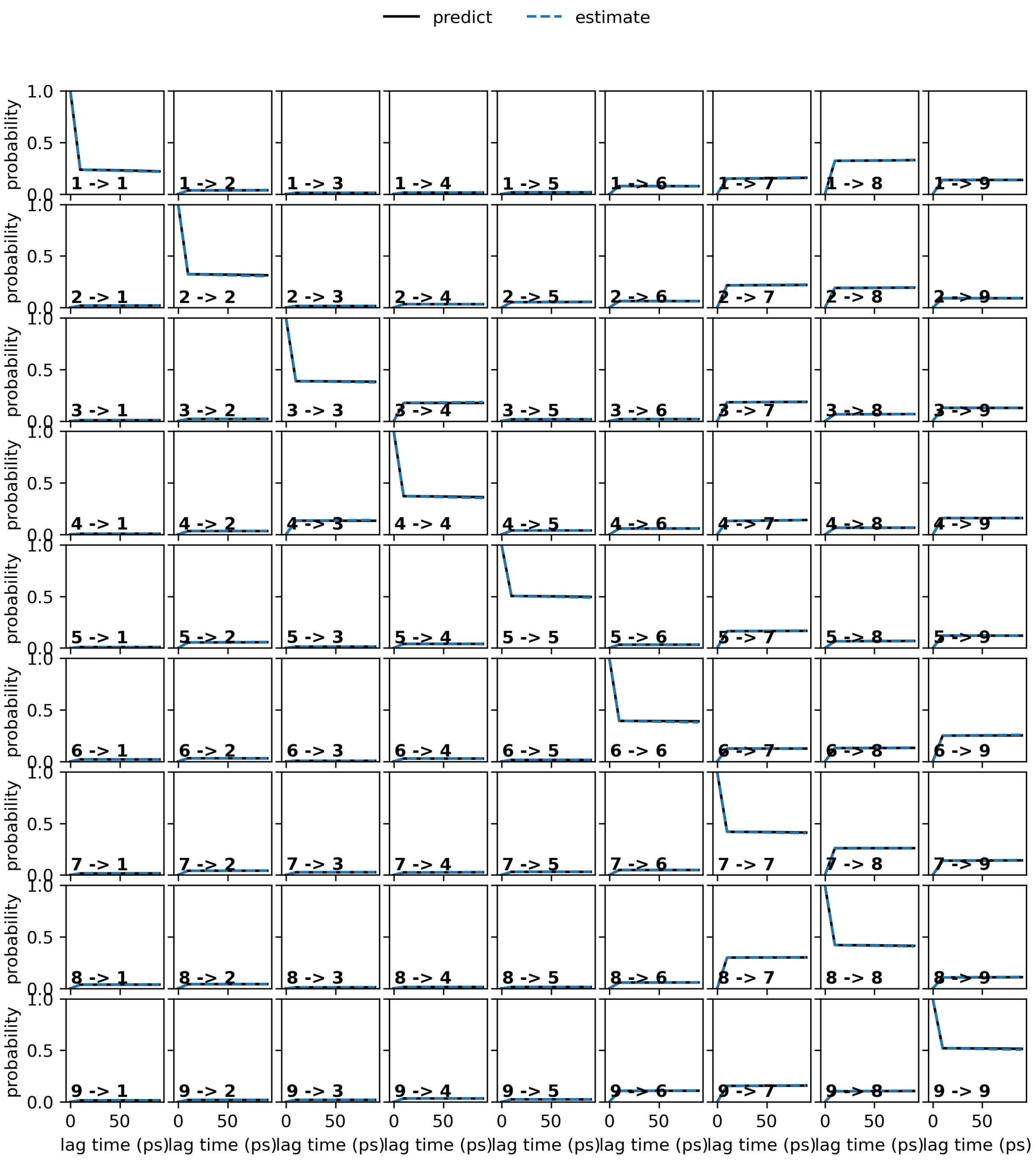


FIG. S2. CK-test for the generated data.
